# Imaging neuronal voltage beyond the scattering limit

**DOI:** 10.1101/2023.12.03.568403

**Authors:** Tsai-Wen Chen, Xian-Bin Huang, Sarah E. Plutkis, Katie L. Holland, Luke D. Lavis, Bei-Jung Lin

**Author notes:** Correspondence (T.-W.C.), (B.-J.L.).

## Abstract

Voltage imaging is a promising technique for high-speed recording of neuronal population activity. However, tissue scattering severely limits its application in dense neuronal populations. Here, we adopted the principle of localization microscopy, a technique that enables super-resolution imaging of single-molecules, to resolve dense neuronal activities *in vivo*. Leveraging the sparse activation of neurons during action potentials (APs), we precisely localize the fluorescence change associated with each AP, creating a super-resolution image of neuronal activities. This approach, termed **A**ctivity **L**ocalization **I**maging (ALI), identifies overlapping neurons and separates their activities with over 10-fold greater precision than what tissue scattering permits. Using ALI, we simultaneously recorded over a hundred densely-labeled CA1 neurons, creating a map of hippocampal theta oscillation at single-cell and single-cycle resolution.

---

Neurons compute and communicate using brief electrical signals known as action potentials (APs). Lasting only about one millisecond, these signals carry rich information about our perception, actions, and thoughts. Modern voltage-sensitive fluorescent proteins (*1–9*) have enabled non-invasive monitoring by converting APs into rapid changes in fluorescence intensity. However, challenges persist in extracting this information from dense neuronal populations *in vivo*.

Imaging voltage signals in densely-packed neurons requires exquisite resolution in both space and time. While two-photon microscopy (*10*) provides excellent spatial resolution, its speed is limited, particularly when trying to image large populations of cells (*5, 7, 11, 12*). On the other hand, wide-field imaging using a high-speed camera allows kilohertz frame rate recordings of neuronal APs (*1–4, 8, 9*). However, tissue scattering makes it difficult to resolve closely-positioned cells, severely limiting the number and density of neurons that can be simultaneously studied (*1–4, 8, 9*).

Here, we introduce Activity Localization Imaging (ALI), a technique that overcomes the resolution limit of scattering, enabling large-scale, kilohertz voltage imaging in dense neuronal population. The underlying principle is akin to localization microscopy in super-resolution optical imaging (*13–15*) (see fig. S1 for comparison). Specifically, densely labeled neurons are difficult to resolve because scattering broadens the distribution of fluorescence, creating ‘footprints’ of individual neurons that overlap. However, at any given moment, only a sparse subset of neurons fires AP and exhibits changes in fluorescence intensity. These changes highlight the active neurons, making them uniquely visible from their neighbors. If one can localize each activation event more precisely than the size of footprints, nearby neurons could be resolved, even when their footprints overlap.

In fact, positional information has been useful for the sorting of APs in extracellular electrophysiological recordings (*16, 17*). Given the high pixel density in imaging experiments, the potential for achieving precise localization offers a promising avenue to resolve neuronal activities beyond the scattering limit.

## Activity localization imaging

We demonstrate ALI using in vivo voltage imaging in mice. Figure 1A illustrates an example of our experiment, in which dense populations of hippocampal neurons expressing the voltage indicator ‘Voltron2’ were simultaneously imaged at 2000 frames per second *in vivo*. The data consist of movies (360,000 frames for a 3-minute recording), in which APs appear as brief (1-2 frames) and small (∼2-6%) fluorescence changes specific to each cell (see Movie S1). Due to scattering, there are often situations where labeling density prevents an unambiguous identification of neurons (Fig. 1A, right).

**Figure 1.**
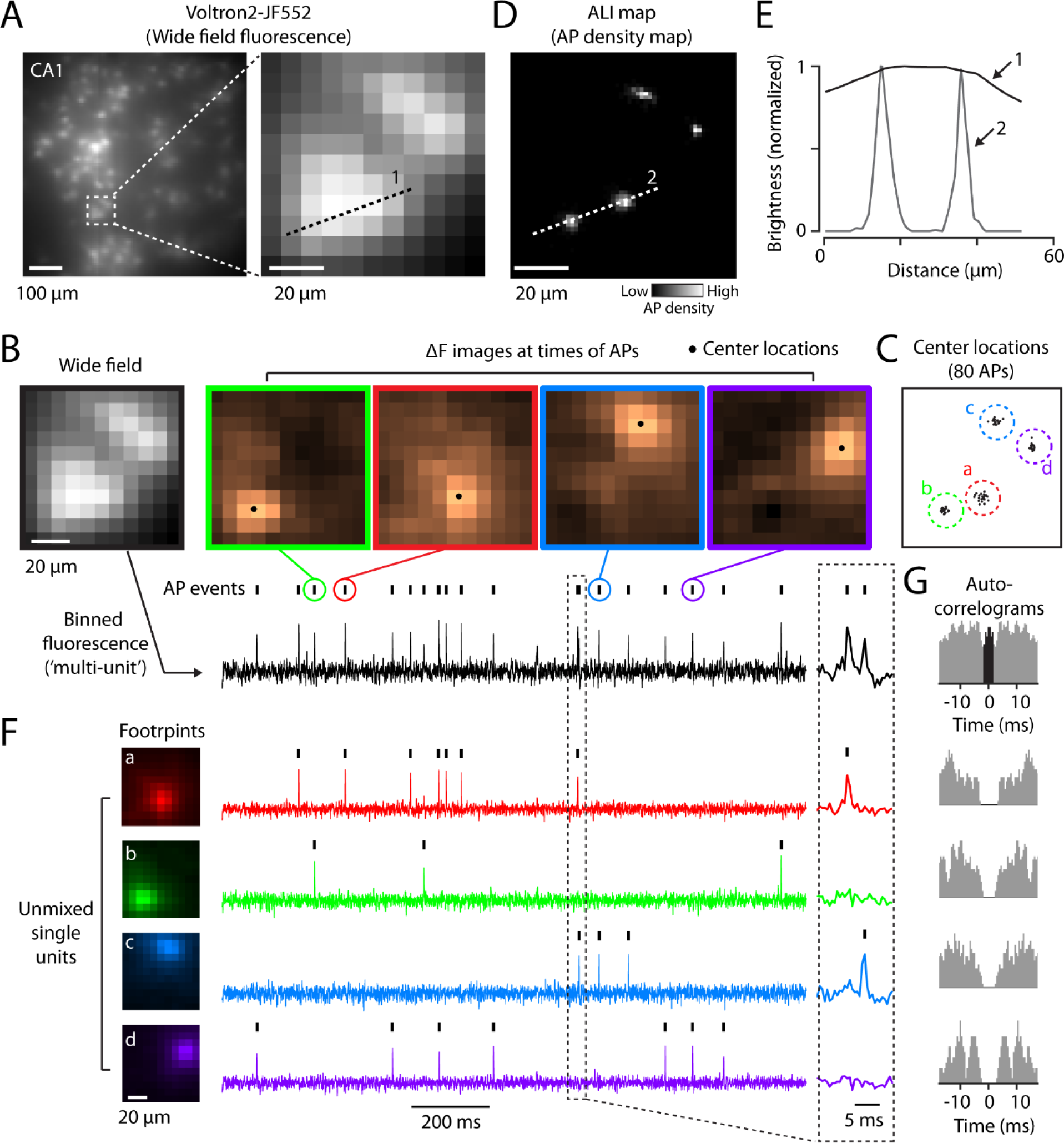
The principle of ALI. (A) Left, a wide-field image of CA1 neurons densely labeled with the voltage indicator Voltron2-JF552. Right, expanded view of the boxed region, showing the difficulty to resolve closely-positioned cells. (B) Densely labeled neurons (the left most image) can be resolved when they are sparsely active during action potentials (APs, indicated by the black ticks above the trace at bottom, representing the binned fluorescence intensity of this region, or ‘multi-unit’ activity. The fluorescence signal was high-pass filtered and inverted to highlight AP events). Each ΔF image (the four images at the right) shows pixel-wise fluorescence change at the peak time of the indicated AP events. The central location of the change (black dots) was determined with sub-pixel precision and overlaid on the respective ΔF image. (C) The central locations of 80 APs plotted on the same reference frame. Four clusters of APs (a-d, marked by different colors) can be distinguished by their distinct central locations. (D) The central locations of all APs detected in this region (n = 1588) shown as a probability density map (i.e., the ‘ALI image’), in which brightness is proportional to the probability of AP occurrence. For clarity, the brightness of each cluster is normalized to its peak. (E) The brightness along lines ‘1’ in wide field and line ‘2’ in the ALI image. (F) Footprints of the four clusters (left) and the corresponding unmixed traces (middle). Boxed region is shown at an expanded time scale (right). (G) Spike auto-correlograms for the multi-unit (top panel) and single-unit (bottom four panels) spike trains. Note the absence of short inter-spike-interval events (i.e., ‘refractory period’) in single-unit but not multi-unit activity.

To better visualize these neurons, we first detect all APs, irrespective of their cellular origins, as peaks in a high-pass filtered fluorescence movie (Fig. 1B, ‘multi-unit’ activity, black ticks indicate detected AP events). We then take the “ΔF image”, a pixel-wise map of fluorescence changes at the peak time of each AP, as signatures to differentiate APs of different cells (Fig. 1B, “ΔF images”). To reduce noise associated with these tiny signals, we performed singular value decomposition-based de-noising to further enhance contrast (see Methods and fig. S2).

Examining these ΔF-AP images reveals distinct activation patterns for different APs, suggesting the presence of multiple neurons. For each ΔF-AP image, we precisely determined the central location of activation, yielding coordinates x0, y0 for the location of each AP (the black dots in Fig. 1B and C, see Methods). Using these coordinates, which we determined for all APs detected in a recording, we generated a spatial map of AP density (Fig. 1D), with brightness proportional to the likelihood of AP occurrence at a specific location.

This map, which we referred to as an ‘ALI’ image, allows the visualization of neuronal activity with a resolution several-fold greater than the limitation imposed by scattering (Fig. 1E). In this example, four distinct clusters of APs are resolvable (Fig. 1C and D), in sharp contrast to the blurry image of wide field fluorescence (Fig. 1A). Within each cluster, AP locations spread over a much smaller area than the typical size of footprints (over 10-fold in some cases, fig. S3), demonstrating our ability to localize APs with high precision. A simple peak-detection algorithm segments these APs into clusters, permitting the separation of activities into distinct, well-isolated sources.

Given the membership information of APs in these clusters, we further determined the underlying footprints (see the colored images on the left of Fig. 1F) by averaging all ΔF-AP images belonging to the same cluster. These footprints reveal the extent to which brain tissue scatters the signal of an individual cell. At a depth of 140 μm, footprints had an average width of 35 ± 9 μm (full width at half maximum), much wider than the typical separation between hippocampal neurons. Furthermore, footprints decay slowly in space, and were much wider in out-of-focus neurons (fig. S4), causing extensive crosstalk affecting large numbers of cells.

Despite this crosstalk, we achieve excellent signal separation with the help of ALI. By identifying overlapping neurons and their footprints, we extract activity traces using least-square regression (colored traces in Fig. 1F, see Methods). These traces demonstrate remarkably low crosstalk. Even for cells that were completely merged into a single spot in the wide field image (e.g., clusters ‘a’ and ‘b’ in this example), individual APs caused little contamination in adjacent neurons (Fig. 1F, inset). Spike auto-correlograms demonstrate a clear refractory period (Fig. 1G), consistent with a low contamination between cells (*18*).

## ALI outperforms existing cell identification algorithms

We compared ALI with VolPy (*19*), a popular analysis pipeline tailored for voltage imaging data (*5, 6*). VolPy identifies neurons using a deep learning model, ‘Mask R-CNN’, comparing favorably with existing algorithms (*6, 19*), on par with human annotators (*19*). We tested ALI and VolPy using simulated data, where ground-truth information is available (Fig 2A). We simulated two neurons with a fixed footprint size (width = w) and number of APs, while systematically increasing the overlap between cells (Fig 2B, the left most column). The separation, s, between neurons was set to s = 1.25w, 1.0w, 0.75w, 0.5w and 0.25w, respectively.

At high separations (s ≥ w), both algorithms successfully identified the two simulated neurons (Fig. 2B, top two rows). However, when the overlap exceeded the scattering limit (i.e., s < w), neuron images began to merge, and VolPy failed to report the presence of two cells in the movie (Fig. 2B, bottom three rows, middle). The extracted trace of VolPy was a mixture of the ground-truth spike trains of the two cells (Fig. 2C, top row).

**Fig. 2.**
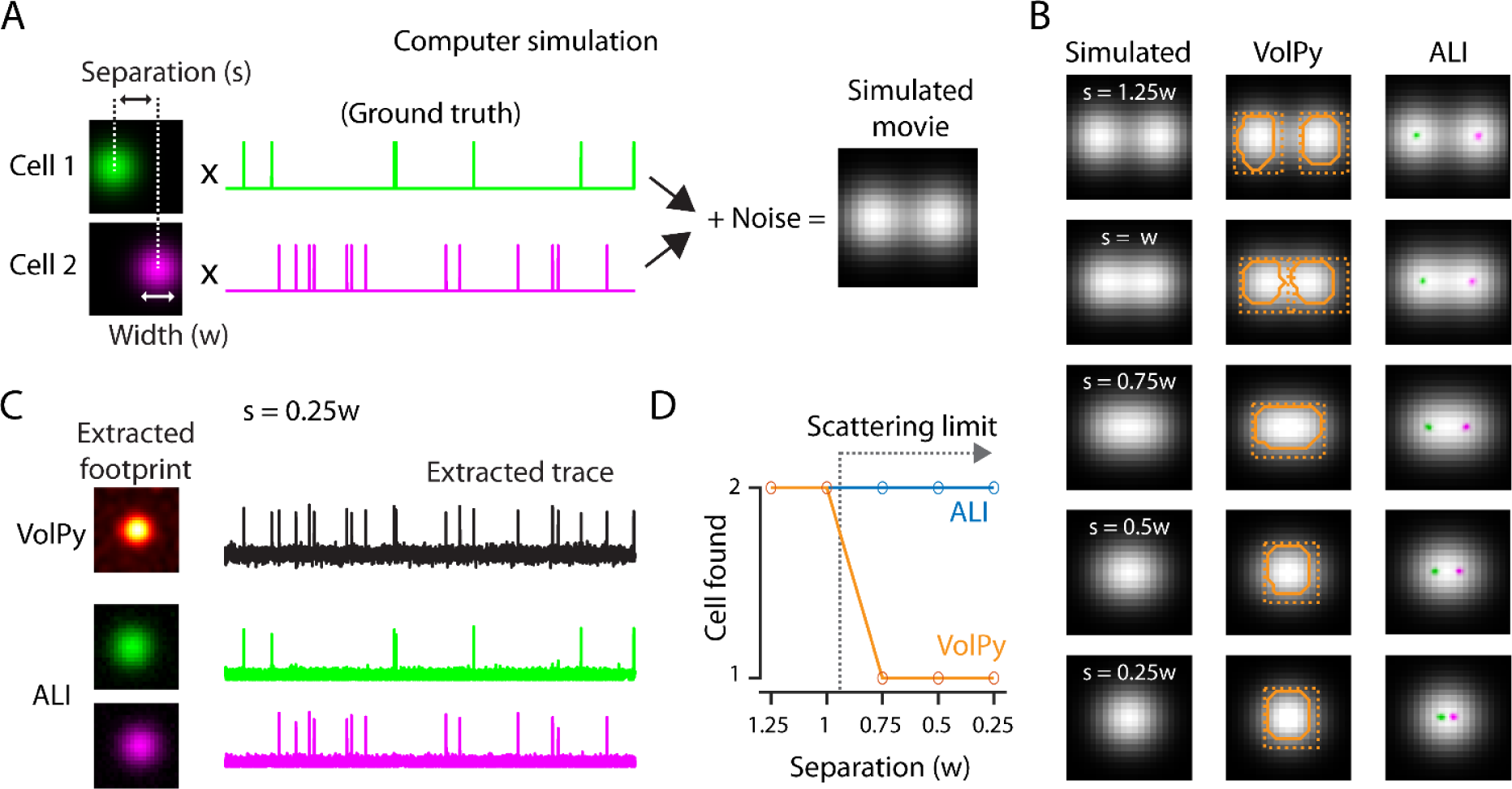
ALI outperforms VolPy in resolving closely-positioned cells. (A) Schematics of computer simulation. Footprints were simulated as 2D-gaussian functions with full width ‘w’ at half the peak intensity. The separation, s, between neurons were varied systematically. Simulated movies were generated by multiplying footprints with ground-truth spike trains, summing across the two neurons, and adding random noise. (B) Mean images of the simulated movies (left), cell identification results of VolPy (middle) and ALI (right). (C) Cell extraction results in highly overlapping scenario (s = 0.25w). The extracted footprints (left) and traces (right) were shown for VolPy (top) and ALI (bottom). (D) The number of neurons found by the two algorithms plotted against the separation between neurons.

In contrast, armed with the precise locations of APs (Fig. 2B, the rightmost column, ALI maps colored and overlaid on the mean image), ALI accurately resolved the two simulated neurons down to the smallest separation (Fig. 2, B to D). It faithfully recovers the footprints of both neurons (Fig. 2C, bottom, two images on the left) and extracts activity traces that mirror the ground-truth spike trains (see Fig. 2C, traces at bottom, vs. ground-truth in Fig. 2A).

## ALI imaging in large populations of neurons

The ability to resolve densely-packed neurons greatly increase the numbers of neurons that can be imaged simultaneously. Fig. 3A shows an example where we simultaneously imaged >100 densely labeled hippocampal neurons at 2000 frames per second.

**Figure 3.**
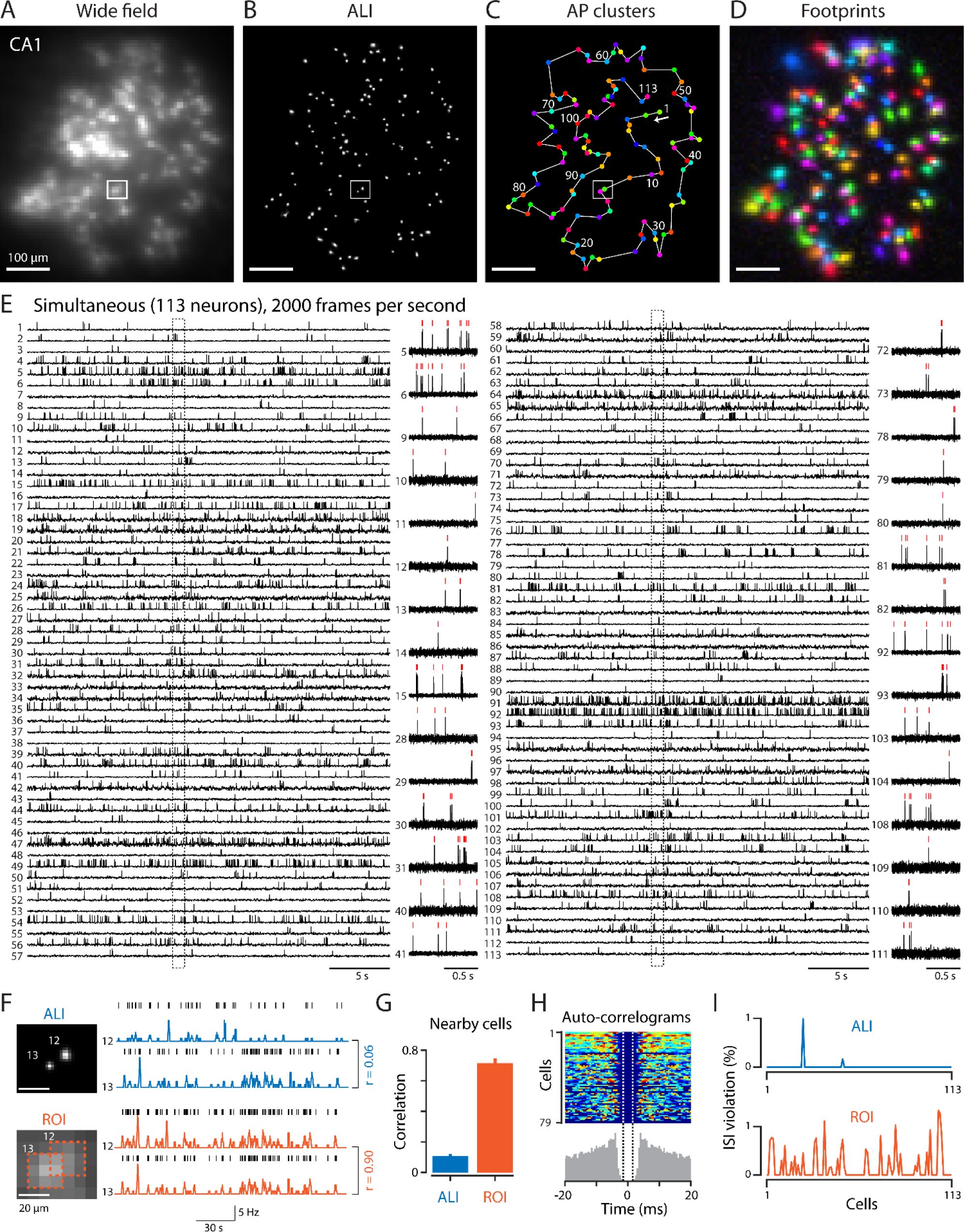
ALI imaging in large neuronal population. (A-D) From left to right: wide field, ALI, AP clusters, and footprints. (E) The extracted signals of the 113 AP clusters, numbered along the white line in (C). Each row shows the extracted trace of one neuron. Boxed areas are expanded in inset. Red ticks indicate detected APs. (F) Left, expanded view of the boxed region in (B) (‘ALI’) and in (A) (‘ROI’). Orange dashed lines indicate regions in which ROI signals were measured. Right, spike trains of cell 12 and 13 (black ticks) and the corresponding spike rate signals in 1s sliding windows (colored traces) shown for ALI (top, blue traces) and ROI (bottom, orange traces), respectively. The numbers on the right indicate correlation coefficients. (G) Correlation (mean ± s.e.m.) between the spike train of each neuron and that of its nearest neighbor. (H) Spike auto-correlograms (ACG) of the ALI spike trains shown as a color-coded image. Each row shows the ACG of one neuron (only cells with more than 100 spikes are shown). The gray histogram at bottom shows the average ACG of all cells. Dashed vertical lines indicate inter-spike-interval (ISI) period in which the occurrence of a spike is counted as an ISI violation event (± 1.5 ms). (I) Percentage of ISIs that are shorter than 1.5 ms (‘ISI violation’).

To apply ALI to such a data set, we first divided images into smaller patches and performed spike detection, de-noising, and localization separately in each patch (see Methods for detail). We then assembled the position information of all APs detected in different patches into a single ALI image. Fig. 3B shows an example of such an image, representing the position information of 23227 APs detected in this recording. These APs were further separated into 113 spatially distinct clusters (Fig. 3C), each containing 206 ± 298 spikes (range 6-1864 spikes per cluster). The membership information of APs in these clusters were then used to determine footprints (Fig. 3D) and the corresponding activity traces of individual cells (Fig. 3E, and fig. S5 and S6).

Examining these traces reveals a dazzling diversity of hippocampal CA1 activity patterns. Nearby neurons, even overlapping ones, exhibit highly distinct spiking patterns. To visualize these patterns, we sorted neurons along the shortest distance path that traverses all neurons exactly once (Fig. 3C, cell #1 to #113 numbered along the white line connecting all neurons). Consecutive traces in 3E thus represent the activity of closely-positioned cells. On average, nearby neurons were separated from each other by 19 ± 10 μm, a distance that falls within the average size of footprints. However, their APs were clearly resolvable, forming distinct clusters in space (e.g., see Fig. 3F, top). The occurrence of AP in one neuron caused little contamination in its neighbors (Fig. 3E, consecutive rows). Nearby spiking patterns exhibit low correlation (Figs. 3F and G, ‘ALI’), even for the most closely-positioned cells (e.g. Fig. 3F, top).

Such distinct dynamics cannot be resolved using traditional analysis methods based on regions of interest (ROIs) selection. To illustrate this, we placed small ROIs (3×3 pixels) at the same locations of the ALI clusters and measured the corresponding spiking patterns by averaging fluorescence within each ROI (see Fig. 3F, bottom). Contrary to the low correlation between ALI-reported activities, activities of nearby ROIs exhibited high correlation (Fig. 3, F and G, ‘ROI’), indicating that these ROIs cannot resolve overlapping neuronal dynamics. Furthermore, while clear refractory periods are visible in the ALI-reported spike trains (Fig. 3H), the ROI spike trains exhibit inter-spike-intervals (ISI) that often fall within the refractory window (‘ISI violation’ events, see Fig. 3I), indicating strong contaminations from nearby cells (*18*).

## ALI clusters correspond to structurally identified neurons

Because ALI resolves many clusters that were not visible in the wide-field image, we ask whether these clusters correspond to structurally identifiable neurons. To address this question, we use a two-photon microscope to image the same field of view after the voltage imaging experiment.

Examining the two-photon z-stack reveals that the clusters identified by ALI indeed correspond to individual, structurally identified neurons (Fig. 4). Overall, all clusters revealed by ALI (113/113) can be mapped to neurons identified in the two-photon z-stack (see Fig. 4C, cells highlighted in yellow). The spatial constellation of these clusters matches precisely with neuron locations in two-photon images (see Fig. 4, B vs. C, boxed regions expanded in D to I). In some cases, two neurons located at slightly different depths were separated in the x-y plane by less than the width of a single neuron (Fig. 4I) and they appeared as a single spot in the wide-field image (Fig. 4G). Even in such cases, ALI separates these closely-positioned APs into distinct, well-isolated clusters (Fig. 4H).

**Figure 4.**
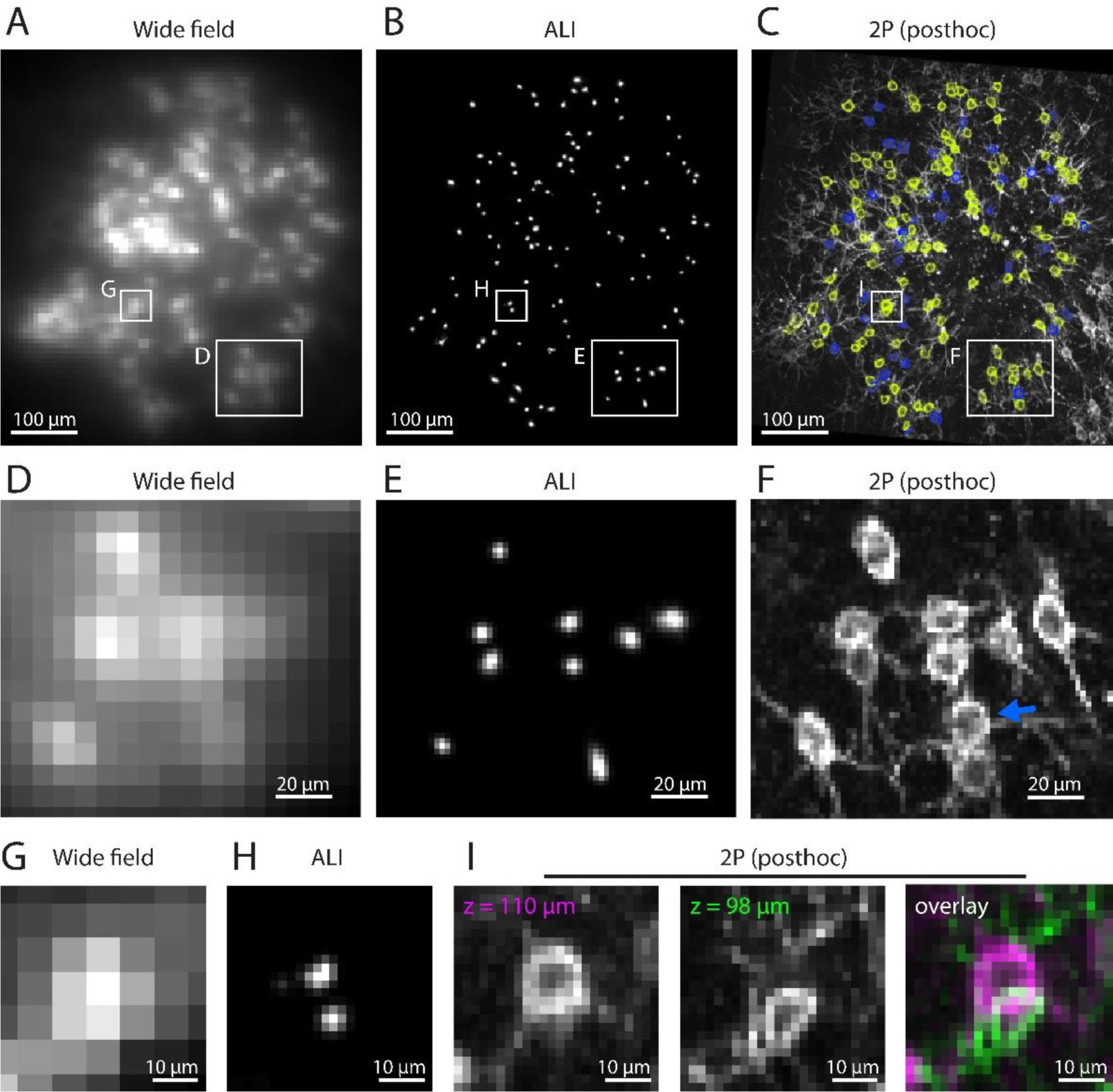
ALI clusters correspond to structurally identified neurons. (A-C) From left to right: wide field (A), ALI (B), and two photon z-stack (C). Neurons corresponding to the ALI clusters are highlighted in yellow (n = 113 cells). Nearby, unidentified neurons are highlighted in blue (n = 44 cells). (D-E) Expanded views of the areas labeled in (A-C). The blue arrow in (F) indicates a neuron that was not found in ALI. (G-I) Expanded views of the areas labeled in (A-C). Two neurons at slightly different z-planes were separated in the x-y plane by less than a cell diameter (I) and were clearly resolved by ALI (H).

Although all clusters revealed by ALI were found in two-photon images, some neurons visible in the two-photon z-stack were missing in the ALI map (see Fig. 4F, blue arrow, and Fig. 4C, cells highlighted in blue, n = 44/157 cells). These neurons may correspond to ‘silent cells’ in CA1 (*20*) that did not fire any AP during the recording period and, therefore, were not visible in the AP density map. Alternatively, AP-related signals may be too weak to be detected, or too closely-positioned to be separated from the APs of an existing cluster. In the latter scenario, these unresolved neurons may contribute to non-zero ISI-violation that we observed in a small number of clusters (e.g. Fig. 3I, top), and could be excluded by setting a minimum threshold for ISI-violation, as is routinely done in electrophysiological recordings (*18*).

## ALI reveals a map of hippocampal theta oscillation

ALI resolves neuronal spiking dynamics with high density, scale, and signal separation. We use these unique advantages to map population spiking dynamics during hippocampal theta oscillations (4-10 Hz), a brain rhythm critical for synaptic plasticity and memory formation (*21*). Neural activities in this frequency range have, until now, eluded analysis through calcium imaging due to insufficient temporal resolution. Although micro-electrode recordings can track spiking dynamics at high frequency (*22*), they lack the density and resolution to map the spatial arrangement of such dynamics, particularly in neighboring cells. Consequently, the fine-scale architecture of neuronal involvement in theta rhythms remains largely unclear.

We use ALI to record large populations of densely-packed CA1 neurons in awake behaving mice (Fig. 5A). Although theta rhythms are typically demonstrated as local field potential (LFP) oscillations, we found that, with >100 CA1 neurons simultaneously recorded, theta-frequency oscillations are directly observable in real-time from population spiking patterns (see Fig. 5B and C, Movie S2). Fig. 5B shows the concurrent spike trains of 110 CA1 neurons, along with their summed activity (‘summed spikes’, bottom), and its theta band filtered version (‘4-10Hz filtered’, top). The population activity exhibits prominent rhythms (see the boxed region of Fig. 5B, expanded below), with a distinct spectral peak at ∼7Hz (Fig. 5C).

**Figure 5.**
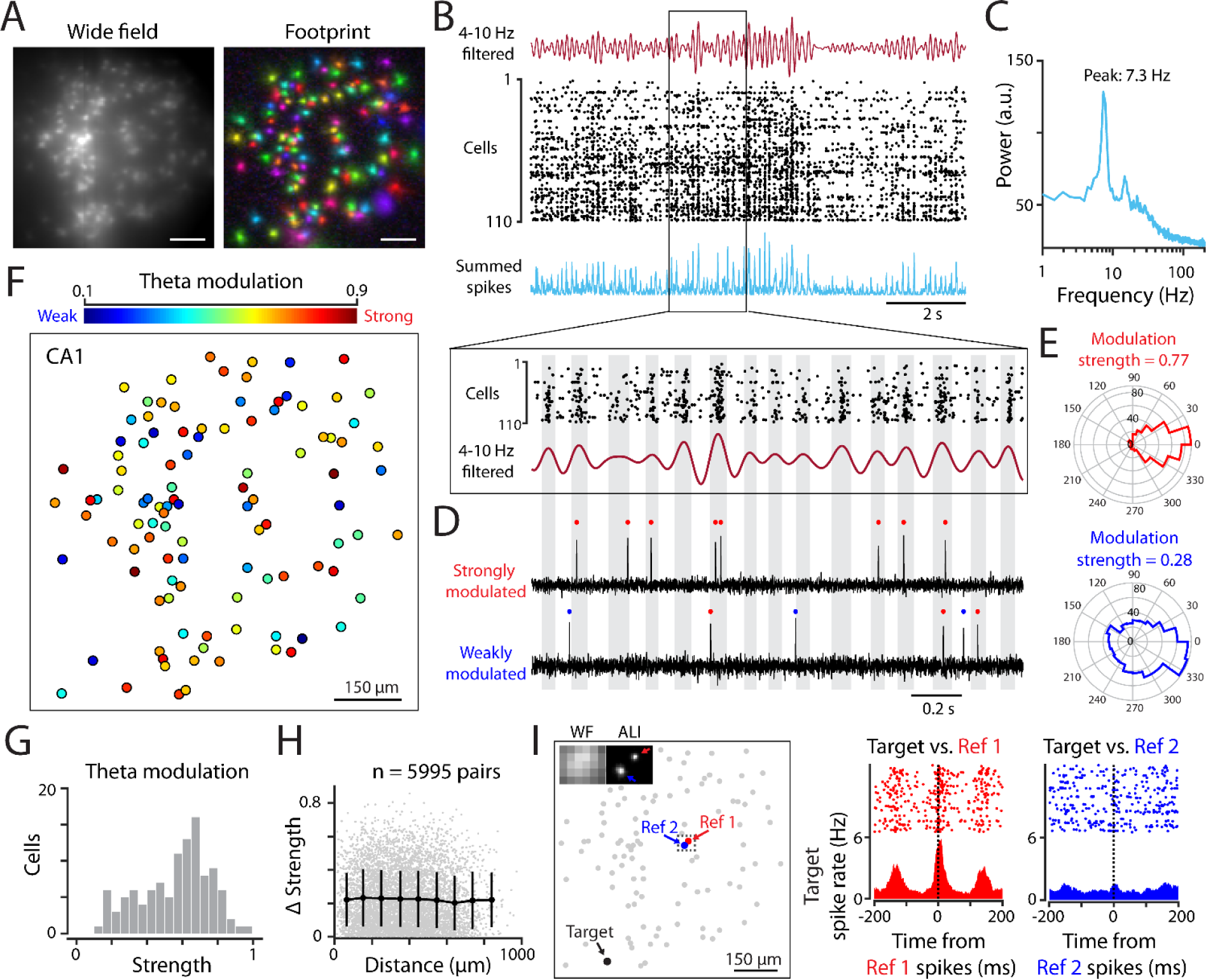
ALI reveals a map of hippocampal theta oscillation. (A) Wide field image (left) and footprints (right) of CA1 neurons. (B) Extracted spike trains of 110 cells (black dots), summed activity across neurons (cyan trace), and its theta-band filtered version (4-10Hz filtered, the trace on the top). The boxed area is expanded below. (C) Fourier spectra of the population spiking activity. (D) Spiking activities of two example neurons relative to the population rhythms. Red and blue dots indicate spikes that occurred during the positive phase (gray shaded areas) or negative phase (white spaces in between) of the oscillation, respectively. (E) Polar histograms of spike phases of the two neurons in (D). The numbers on top show their respective theta modulation strengths. (F) A color-coded map showing the spatial distribution of theta modulation strength. Each circle shows a neuron. Color indicates the strength of theta modulation (blue to red: weak to strong). (G) A histogram of theta modulation strength for all neurons. (H) Pairwise difference in modulation strengths plotted against the distance between neurons (each dot indicates a pair of cells, n = 5995 pairs). The black curve shows mean ± SD in each distance bin. (I) Left, location of three example neurons (Target, Ref1, and Ref2). Inset: corresponding wide field (WF) and ALI image of the boxed region. Right, spike cross-correlograms of two cell pairs (‘Target’ vs. ‘Ref 1’ and ‘Target’ vs. ‘Ref 2’) showing their varying degree of theta coherence.

We analyzed how individual neurons contribute to this population rhythm. Consistent with previous studies (*22*), individual CA1 pyramidal neurons typically spike sparsely, contributing either zero or a few spikes in each oscillation cycle (Fig. 5D). The degree of phase modulation varies greatly from cell to cell: while some neurons spiked in a highly specific phase of the population rhythm (‘strongly modulated’ cells, Fig. 5D and E, top), other neurons showed a much broader distribution in spike-phases (‘weakly modulated’ cell, see Fig. 5D and E, bottom). For each neuron, we quantified the strength of theta modulation using an index ranging from 0 (no modulation) to 1 (strongly modulated, see Methods). This index exhibits a broad distribution (Fig. 5F and G), with different neurons showing a wide range of modulation strengths.

ALI allows us to map the spatial arrangement of this modulation strength at the cellular level. Fig. 5F shows an example of such a map, in which different modulation strengths are shown by different colors (blue and red colors represent weak and strong modulation index, respectively). Examining this map reveals a fine-grained spatial arrangement of theta modulation: nearby cells, even the adjacent ones, often exhibit highly distinct modulation strengths (Fig. 5F). Pairwise difference in modulation strengths was as high in nearby neurons as in more distant cell pairs (Fig. 5H).

In addition to the heterogeneous modulation strength, we found substantial variations in the pairwise coherence between neurons (Fig. 5I and fig. S7). For example, a neuron (‘Ref 1’) can exhibit strong coherence with a specific target cell (‘Target’), while its immediate neighbor (‘Ref 2’) does not (see spike correlograms in Fig. 5I, right). These neurons were highly overlapping in space, and their APs were separable only based on central locations (Fig. 5I, inset), yet they exhibit clearly distinct functions. These data reveal remarkable specificity in theta-frequency communication between hippocampal neurons. Furthermore, they highlight the power of ALI in resolving the detailed functional arrangements in densely overlapping cells.

## Discussion

By precisely localizing APs, we present a simple approach to resolve dense neuronal activity in scattering tissues. This establishes precise localization as a general approach to improve image resolution, even when light diffraction (*13, 14*) is not the primary constraint. In vivo, ALI resolves neurons beyond the capability of human observers, distinguishing it from existing cell finding methods (*19, 23–29*). As we demonstrate here, >100 densely labeled neurons can be simultaneously imaged at high speed and signal separation, providing a density and scale of recording unmatched by existing techniques.

These recordings allow for a direct observation of oscillatory structures in population spike trains, without reference to field potentials. Furthermore, specific communication can be discerned in neurons with highly overlapping footprints. Importantly, individual spike trains can be mapped to structurally identified neurons in anatomical images. This could facilitate subsequent analyses that relate *in vivo* spike trains to circuit connectivity (*30*), morphology (*31*), and molecular markers of cell types (*32*) during *in vitro* studies.

A key advantage of ALI is that it achieves high resolution not by covering neurons with large numbers of pixels, but by precisely localizing APs. As we demonstrate here, covering a neuron with as few as ∼10 pixels (i.e., ∼3 by 3 pixels per neuron) affords precise localization several-fold beyond single-pixel resolution. This makes it possible to use the limited number of camera pixels for a larger field of view, increasing the number of neurons that can be simultaneously studied. Reducing pixel count also decreases data size and the computational cost of image analysis.

Although we demonstrate ALI for somatic voltage imaging, the same principle might be applicable to resolve other types of spatially overlapping signals. This could include different forms of neuronal functional imaging, such as calcium (*33*) or neurotransmitter imaging (*34*), and perhaps other examples of spatially overlapping signals that are sparse in time.

As a method that identifies neurons by their activities, ALI cannot reveal cells that do not fire any AP during the recording period. In our study, about one third of Voltron-expressing neurons were not visible in the ALI image, consistent with the presence of silent cells in CA1 (*20*). To reveal these neurons, activity could be induced using sensory or optogenetic stimulation, which may allow for a more complete mapping of the entire labeled population.

Going forward, ALI could benefit from improvements in the brightness and sensitivity of available GEVIs, permitting better localization in challenging conditions (*35*). Improvements in photo-stability and recording duration (*5, 8*) will provide more APs to aid the identification of weakly active cells. Algorithmically, including additional features of ΔF images, such as the width or shape of a neuron’s footprint, may further improve separation, possibly along the axial direction (*36*). In terms of imaging hardware, efforts to increase the speed of optical-sectioning microscopes (*11, 12, 37*) will reduce out-of-focus contamination and further improve recording quality. In this respect, the ability of ALI to resolve neurons at low pixel counts will be helpful, as fewer pixels translate to faster imaging speed in scanning microscopes. Together, these efforts will take us ever closer to realizing a complete mapping of voltage dynamics in dense neuronal circuit.

## Methods

### ALI analysis pipeline

A summary of the ALI pipeline is shown in fig. S2. All analysis routines are implemented in MATLAB. Details of each step are described below.

### Pre-processing

Brain movement was corrected by aligning each image frame to a reference image (the mean of the first 2000 frames of the recording). For each frame, we first estimated the amount of movement relative to the reference. This movement was then corrected by shifting each image by the opposite amount through gridded interpolation (using the ‘griddedInterpolant’ function in MATLAB). Parallel computing toolbox in MATLAB was used to improve the speed of computation.

After movement correction, we performed pixelwise temporal high-pass filtering to remove slower fluctuations, isolating AP-specific signals. We do this by first computing a slow baseline trace through temporal median filtering of each pixel’s signal (10 ms time windows). This baseline trace was then subtracted from the signal of each pixel, leaving the fast, AP-specific component. Because the fluorescence of Voltron decreases with membrane voltage, we invert the difference after median subtraction, such that action potentials (APs) appear as positive intensity peaks in the resulting ΔF movie.

If the recording covers a small field of view, subsequent detection, denoising, and localization of AP can be performed directly using the movie of ΔF images as input. However, if the recording covers a large field of view with hundreds of neurons, we find it helpful to first divide the ΔF movie into smaller patches, and perform these steps separately for each patch. In our case, we divide our 160 × 190-pixel movies into rectangular patches of 27 × 27 pixels, with a 14-pixel overlap between neighboring patches.

### AP localization

Given the ΔF movie of each patch, we detect candidate action potentials (APs) as peaks of the ΔF movie exceeding a pre-defined threshold (5x the standard deviation of baseline noise). This step aims at detecting all APs, without distinguishing which APs come from which cell. Subsequently, we take the “spatial waveform” of each AP (i.e., the ΔF image at the peak time of AP, or the “ΔF-AP images”), which contains information to identify APs of different cells.

After detecting APs and collecting corresponding ΔF-AP images, we reduce noise in these images using singular value decomposition (SVD). SVD is a popular dimensional reduction technique that represents each image by a set of orthonormal bases (called the ‘principal components’) that capture its key features. By performing SVD on the set of ΔF-AP images and keeping only the first N principle components (N << the number of pixels), the higher order noises in the ΔF-AP images are effectively suppressed. Although keeping a smaller number of components (i.e., smaller N) can lead to better noise reduction, it is important to keep N larger than the potential number of neurons within the patch so that the distinct footprints of these neurons can be properly represented. In practice, we typically use N = 25 given our patch size and labeling density. For smaller patches or a sparser labeling, an even smaller N value may be used to further reduce noise.

After noise reduction, active neurons become clearly visible in each ΔF-AP image. To determine the center location of these APs, we first identify the brightest pixel in each ΔF-AP image. We then refine the estimate beyond the precision of a single pixel by computing the center of mass of pixels surrounding the peak, as

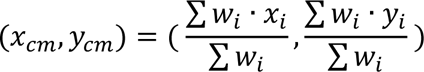

Here, *x*_*i*_ and *y*_*i*_ represent the x and y coordinate of each pixel i, and *w*_*i*_ represents the weight factor, chosen as the square of the pixel’s ΔF value. The summation takes place over the 15 brightest pixels around the peak.

We repeat this procedure to determine the center coordinates of all APs detected in different patches. Because there was overlap between patches, the same AP could be detected more than once by neighboring patches. We therefore exclude an AP if its central coordinate is closer to the center of a nearby patch to avoid duplicate detection.

### Cell finding

With the central coordinates of all APs, we next perform clustering by grouping APs with similar spatial locations. To do this, we first created a 2-D histogram of AP locations (i.e., the ‘ALI’ image) by counting the number of APs in each spatial bin. Because we determine AP locations more precisely than the size of an image pixel, we use a bin size that is 4 times smaller, in both X and Y directions, than the image pixel size to create a “super-resolution” image of AP location. The resulting histogram counts were then smoothed by spatial low-pass filtering, using a gaussian kernel with standard deviation equals to 0.7 times the bin size.

We next identify significant peaks in the smoothed histogram as centers of the AP clusters. We use the following criteria to automatically detect the most prominent clusters. Specifically, for a bin to be detected as a cluster center, it must (i) have the largest count in a spatial neighborhood (radius = 3 bins) (ii) its smoothed count must exceed 4. This approach automatically detected most of the clusters reported in this study (∼90%). However, it can miss weakly active neurons due to a small number of APs, or less well-localized neurons due to broader distributions of AP locations. Both situations result in smaller and noisier peaks in the 2D histogram, making it more difficult to detect these clusters automatically. In such cases, we examined a plot of AP locations (similar to the one shown Fig. 1C) and manually selected these less prominent clusters. Finally, with all clusters identified, individual APs are assigned to the cluster to which they are most closely located.

### Extraction of footprints and voltage traces

With the membership information of APs of each cluster, we next estimate the footprint of by averaging all ΔF-AP images belonging to the same cluster. To ensure footprint purity, we exclude APs that occur synchronously with the APs of other clusters, to prevent images of these synchronous neurons from appearing in the average.

After obtaining the footprints of all clusters, we performed least-square regression to extract the time dependent voltage traces. We first limit the spatial support of each footprint to a circular region around its peak (radius: 10 pixels, or 75 μm), by setting footprint values to zero beyond this region (*38*). We then model the ΔF movie as a linear combination of the footprints, *footprint*_*i*_(*x*, *y*), with *c*_*i*_(*t*) being the coefficient of linear combination for the i^th^ footprint.

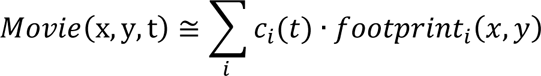

We then determine the coefficient *c*_*i*_(*t*) given the high-pass movie and the set of footprints using least square regression. Because the spike trains are non-negative, a non-negative constraint on *c*_*i*_(*t*) was further imposed to improve performance (*39, 40*).

Given the extracted traces, we refine spike detection using thresholding methods described in (*41*). The refined spike times were used to further improve footprint estimation and trace extraction through the same procedures described above. Since the initial footprint estimation is typically quite good already, this iterative procedure converges quickly (within 1-2 iterations). In practice, we run the iterative procedure twice before settling on a final activity trace, footprint, and spike times of each neuron.

### Computer simulations

To compare the performance of VolPy and ALI, we simulate fluorescence movies (21 pixels × 21 pixels × 80000 frames) consisting of two neurons with varying degrees of overlap. First, we simulate footprints as 2D gaussian functions with full width (w) at half the peak intensity of 8 pixels. The separation, s, between neurons was varied systematically from s = 10, 8, 6, 4, to 2 pixels, corresponding to 1.25w, 1w, 0.75w, 0.5w, and 0.25w. For each neuron, 200 action potentials (APs) were generated at randomly chosen time points. The fluorescence signal was simulated as a constant value that decreases by 10% at the simulated spike times to match the negative polarity of the Voltron indicator. We then multiply the footprint of each cell with this fluorescence signal, summing the resulting movies of both neurons, and adding gaussian random noise to produce the simulated movie. This procedure was performed in MATLAB, and the simulated movies were saved as .h5 files for subsequent analysis using the VolPy pipeline (https://github.com/caichangjia/VolPy).

### In vivo imaging experiments

#### Preparing animals for voltage imaging

All procedures were performed in accordance with protocols approved by National Yang Ming Chiao Tung University Animal Care and Use Committee. Adult C57BL/6 mice of either sexes aged 2 to 3 months were used. To prepare animals for voltage imaging, mice were anesthetized with 1-2 % isoflurane and placed on a heat-pad. A circular craniotomy (3 mm diameter) was performed in the left hemisphere, centered at 2.0 mm caudal and 2.0 mm lateral from bregma. The dura was carefully removed, and the surface of CA1 was exposed by gently removing the overlying cortex with aspiration. A cre-dependent, soma-targeted version of the Voltron 2 virus (AAV-syn-Flex-Voltron2-SOM2) was diluted to 2×10^13^ GC/ml in PBS and injected at four locations at a depth of 200 μm from the CA1 surface. The injection sites were separated from each other by 700 μm. A total 50 nl of viral solution was injected per site at a rate of 1 nl/s. After virus injection, an imaging window consisting of a 3mm diameter cover glass glued to a stainless-steel cannula (3 mm diameter, 1.5mm height) was placed onto the hippocampus and secured to the skull using super-bond C&B (Sun Medical). A titanium head bar was also glued to the skull for head fixation during imaging.

Due to the lack of axial resolution of one-photon voltage imaging, labeled neurons located above or below the imaging plane will contribute to out-of-focus background contamination. To minimize this, we restrict Voltron expression to a thin layer of CA1 neurons. This is achieved by delivering CRE at a specific embryonic day (E14.5 or E17.5) through the injection of AAV-CaMKII-Cre virus (Addgene: 105558-AAV1) into the left lateral ventricle of the animals. This approach labels a specific sub-layer of CA1 neurons (*42*), minimizing the contamination from out-of-focus cells.

#### Voltage imaging

Imaging experiments started 3 weeks after the window surgery. For fluorescence labeling, 100 nanomoles of JF552-Halotag ligand were dissolved in 20μl of DMSO (Sigma), diluted first in 20 μl of Pluronic™ F-127 (20% w/v in DMSO, P3000MP, Invitrogen), and subsequently in 80 μl of PBS. The dye solution was then administered to the animals through retro-orbital injection using a 30-gauge needle. Imaging sessions begin at around three hours post-dye injection.

A 532 nm laser (Opus 532, Laser Quantum) passing through an excitation filter (FF02-520-28, Semrock) was expanded and projected to the sample to illuminate Voltron-expressing neurons. The illumination intensity at the sample plane was ∼70 mW/mm^2^. Fluorescence emitted by the labeled neurons was collected using a 16X, 0.8 NA objective (Nikon), separated from the excitation light using a dichroic mirror (540lpxr, Chroma), an emission filter (FF01-596/83, Semrock), and imaged onto a high-speed CMOS camera (DaVinci-1K, RedShirt) using a 50mm camera lens as the tube lens (Nikkor 50mm f1.2, Nikon). Images were captured at a rate of 2000 Hz using the DaVinci-1K CMOS camera and Turbo-SM64 software (SciMeasure and RedShirt Imaging) for three minutes (360,000 images). Each image had dimensions of 190 × 160 pixels, corresponding to an area of ∼1.4 × 1.2 mm (pixel size was around 7.5 μm). During imaging, animals were fully awake and had the ability to move their whiskers and limbs while being head fixed.

#### Two-photon imaging

After the voltage imaging experiments, animals were removed from the imaging setup and placed under a two-photon microscope. The same area of the CA1 was identified based on blood vessel patterns, and a two-photon z-stack (100 images at increasing depths) was collected at 2-micrometer spacing with an image resolution of 512 × 512 pixels (covering a 580 μm × 580 μm × 200 μm volume of CA1). To excite JF-552 fluorescence, neurons were illuminated using a Ti-Sappire laser (Coherent, Chameleon Ultra II) at a wavelength of 850 nm. Voltron-JF552 fluorescence was collected using a 16X, 0.8 NA objective (Nikon) and separated from the laser light using an emission filter (FF01-609/57-25, Semrock).

#### Analysis of theta modulation

To reveal oscillations in population spiking activity, we first calculate summed activity by counting the number of spikes across the entire recorded population in each 0.5-millisecond time bins. This summed activity trace was then cut in 2-second segments, and the spectral content in each segment was analyzed using a 4096-point fast Fourier transform. The absolute value of the Fourier coefficients was then averaged across segments, revealing the spectral content of the population activity.

To compute theta frequency modulation of individual neurons relative to the population rhythm, we first filter the summed activity trace using a 10^th^ -order Butterworth bandpass filter (passband: 4-10 Hz). The ‘filtfilt’ function in MATLAB was used to ensure zero-phase distortion. The instantaneous phase φ(t) and amplitude A(t) of the filtered trace was determined using Hilbert transform. To quantify spike phase modulation, a vector *V*_*k*_ = *A*(*t*_*k*_)*e*^*i*φ(*t*^_*k*_^)^ was assigned for each spike occurring at time *t*_*k*_. The strength of phase modulation was computed as the normalized vector length:

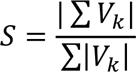

Here, the summation takes place over all spikes of a neuron, and | · | denotes the absolute value of complex numbers.

## Acknowledgments

We thank Drs. Eric Schreiter, Ilya Kolb, and Ahmed Abdelfattah for sharing the Voltron 2 virus, Ms. Hui-Ching Chen for technical assistance, and Dr. Shih-Chieh Lin for comments on the manuscript. During the preparation of this work the authors used ChatGPT 3.5 to improve language and readability.

## Funding

National Science and Technology Council, Taiwan (T.-W.C and B.-J.L)

Howard Hughes Medical Institute, USA (LDL)

## Author contributions

Conceptualization: TWC, BJL

Mouse experiments: XBH, BJL

Analysis/visualization: TWC JF

dye synthesis: SEP, KLH, LDL

Administration/supervision/funding acquisition: TWC, BJL, LDL

Writing: TWC, BJL

---

**Fig. S1.**
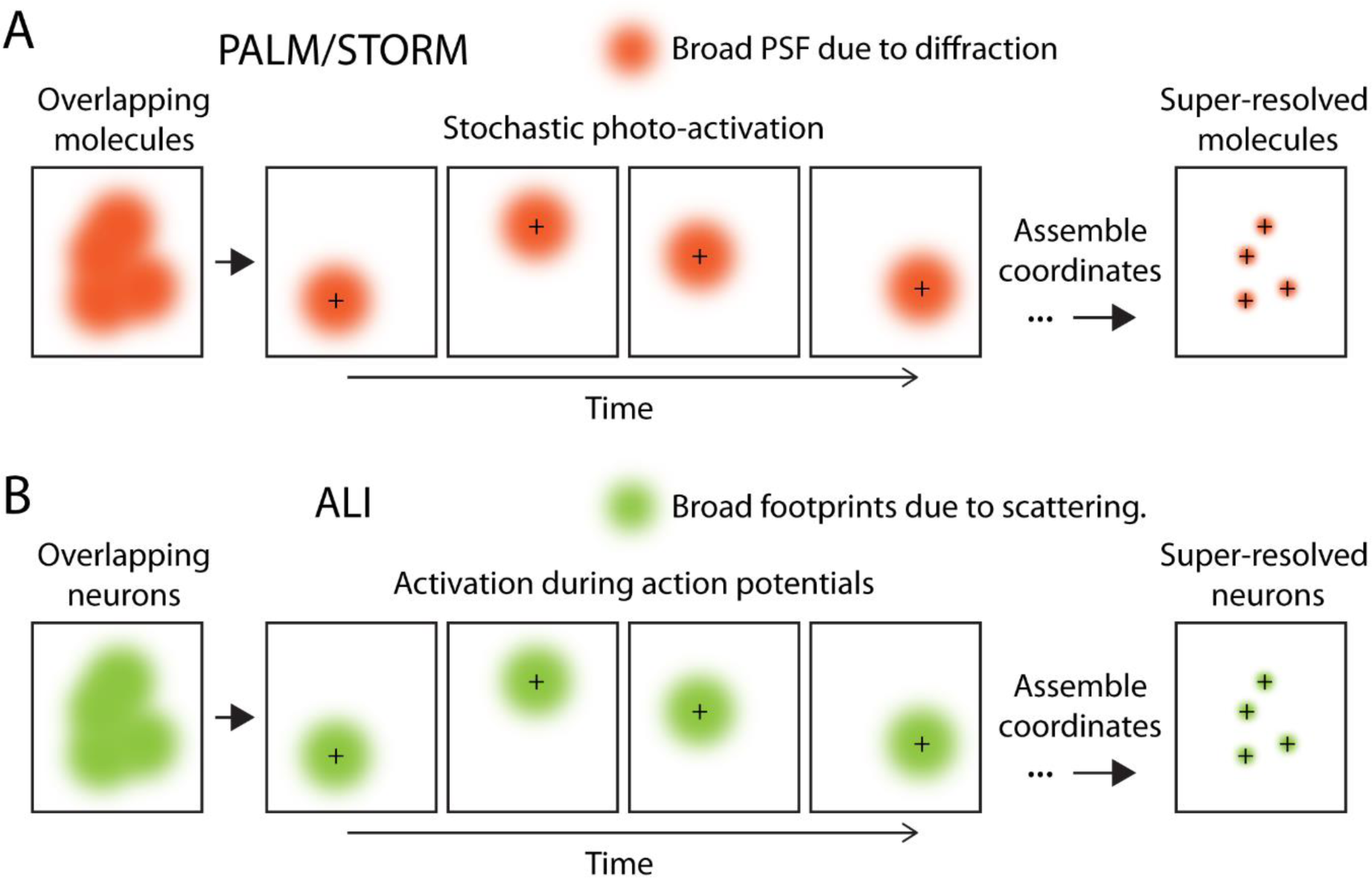
Comparison between localization microscopy techniques and ALI. (A) Localization microscopy techniques, such as PALM and STORM microscopy, aim to overcome the effect of light diffraction, which limits resolution by producing a broad point-spread-function of microscopes (PSF). In these techniques, overlapping molecules (left) are first separated from one another through stochastic photoactivation, which highlights a sparse subset of molecules and permitting their unique visualization (middle). The central coordinate of each active molecule (depicted by the ‘+’ signs) is then determined with high precision and assembled into a super-resolution image (right). (B) By contrast, ALI aims to overcome the scattering of light by brain tissue, which limits resolution by creating a broad ‘footprint’ for each neuron. Here, overlapping neurons (left) are visualized individually when they fire action potentials, which generate specific changes in fluorescence intensity that distinguish the active cell from its neighbors (middle). The central coordinate of each activation event (depicted by the ‘+’ signs) is then determined with high precision and assembled into a super-resolution image, which we termed the ALI map (right).

**Fig. S2.**
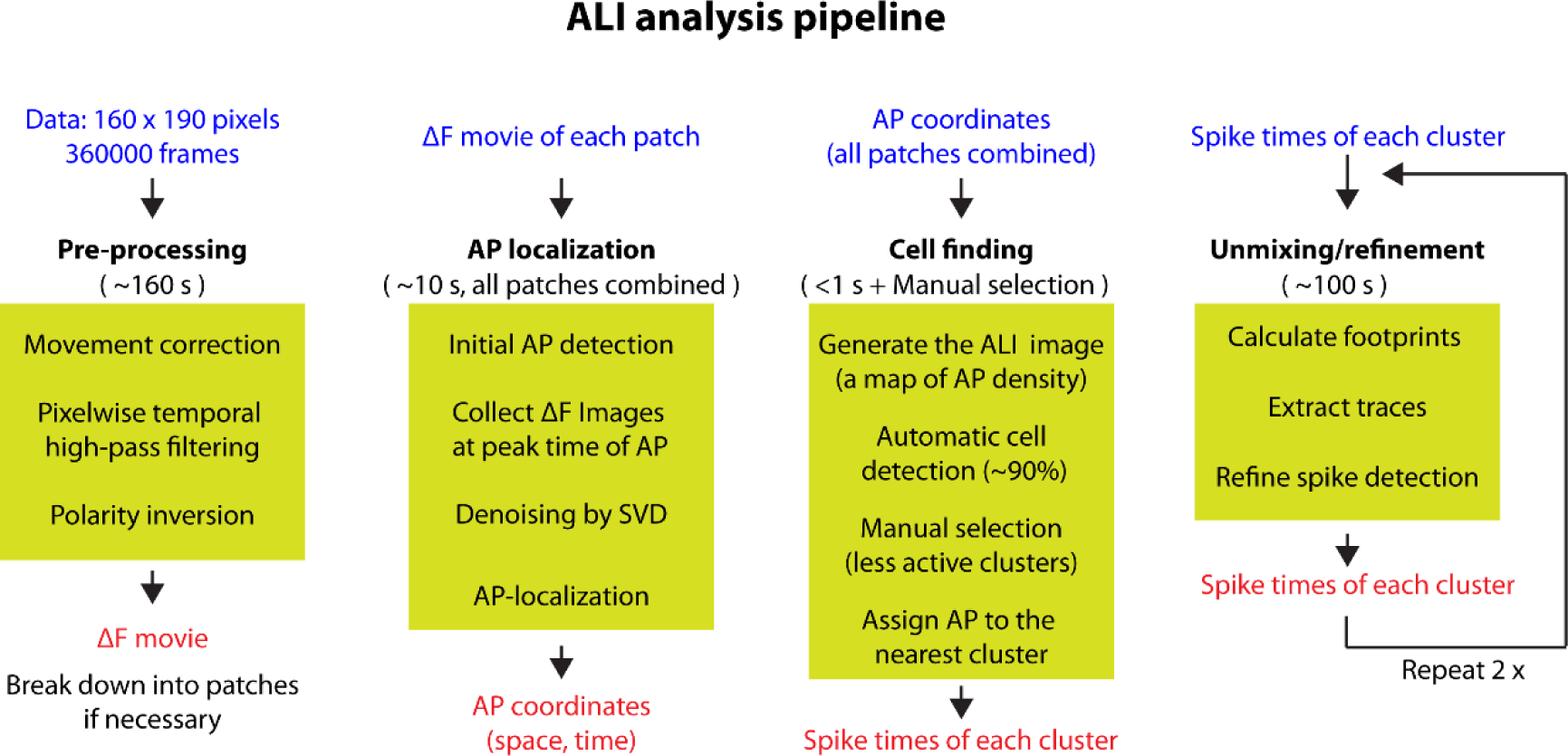
Schematic of ALI pipeline. From left to right: four modules of the ALI analyses pipeline (see Methods for details). Blue and red describes the input and output of each module, respectively. Numbers in the parentheses show the typical computational time of a 160 pixel × 190 pixel × 360,000 frame movie, measured on a desktop PC equipped with a 12th Gen Intel(R) Core i7-12700K 3.61 GHz CPU and 64 GB RAM.

**Fig. S3.**
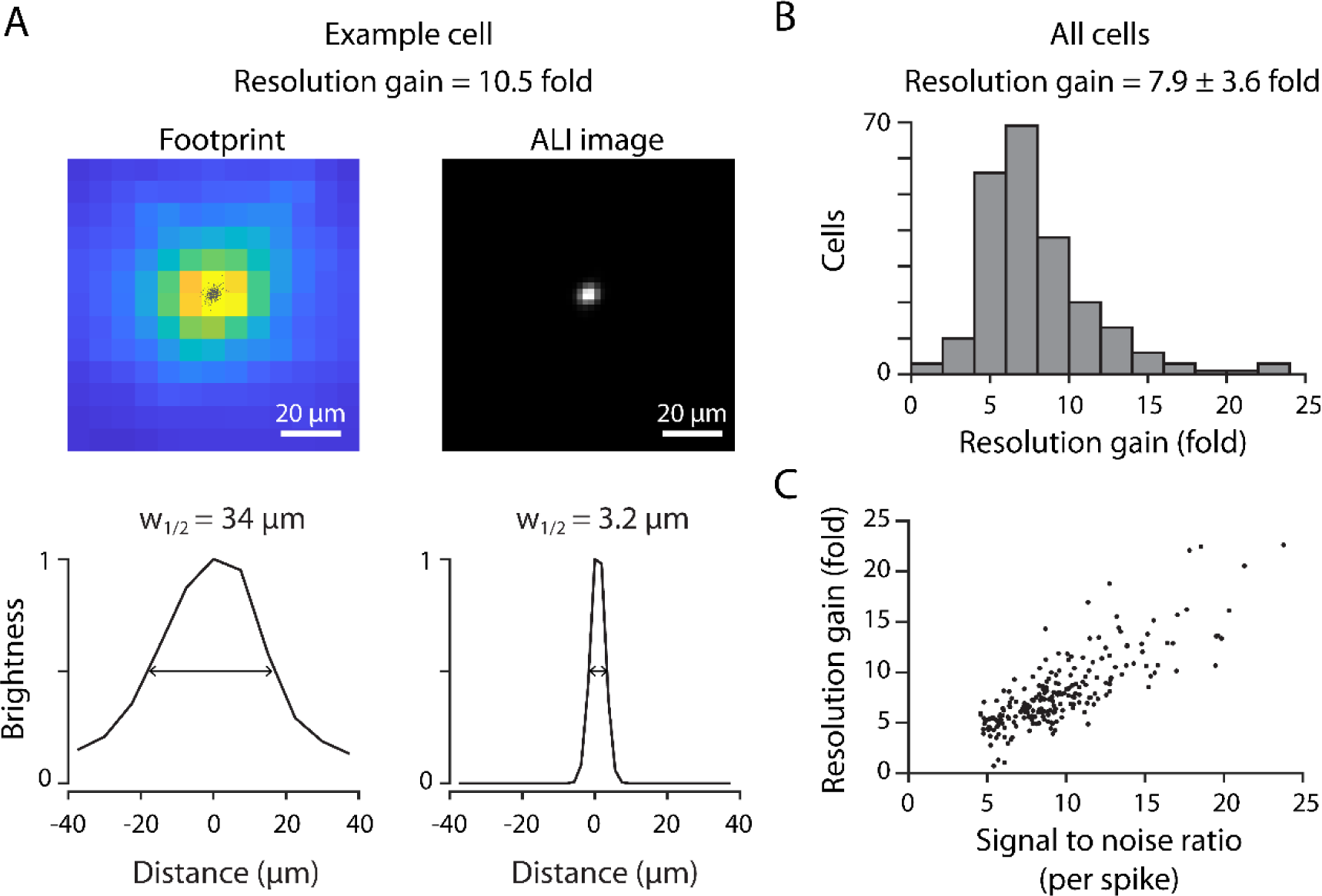
Resolution gain of ALI. (A) Left, the footprint of an example neuron. Black dots overlaid on the image show the central coordinates of individual AP events (n = 323). Right, a 2D density map of AP locations (i.e., ALI image). The curves below show normalized brightness vs. distance along a line passing through the center of the respective images. The numbers indicate full widths at 50% the peak intensity (w_1/2_). The resolution gain is the ratio between the two widths. (B) Resolution gains of 223 CA1 neurons shown as a histogram. The average gain in resolution was 7.9-fold. (C) Resolution gain plotted against the signal-to-noise ratio (SNR) of action potentials. Each dot shows a neuron. Better SNR leads to improved localization and a higher gain in resolution.

**Fig. S4.**
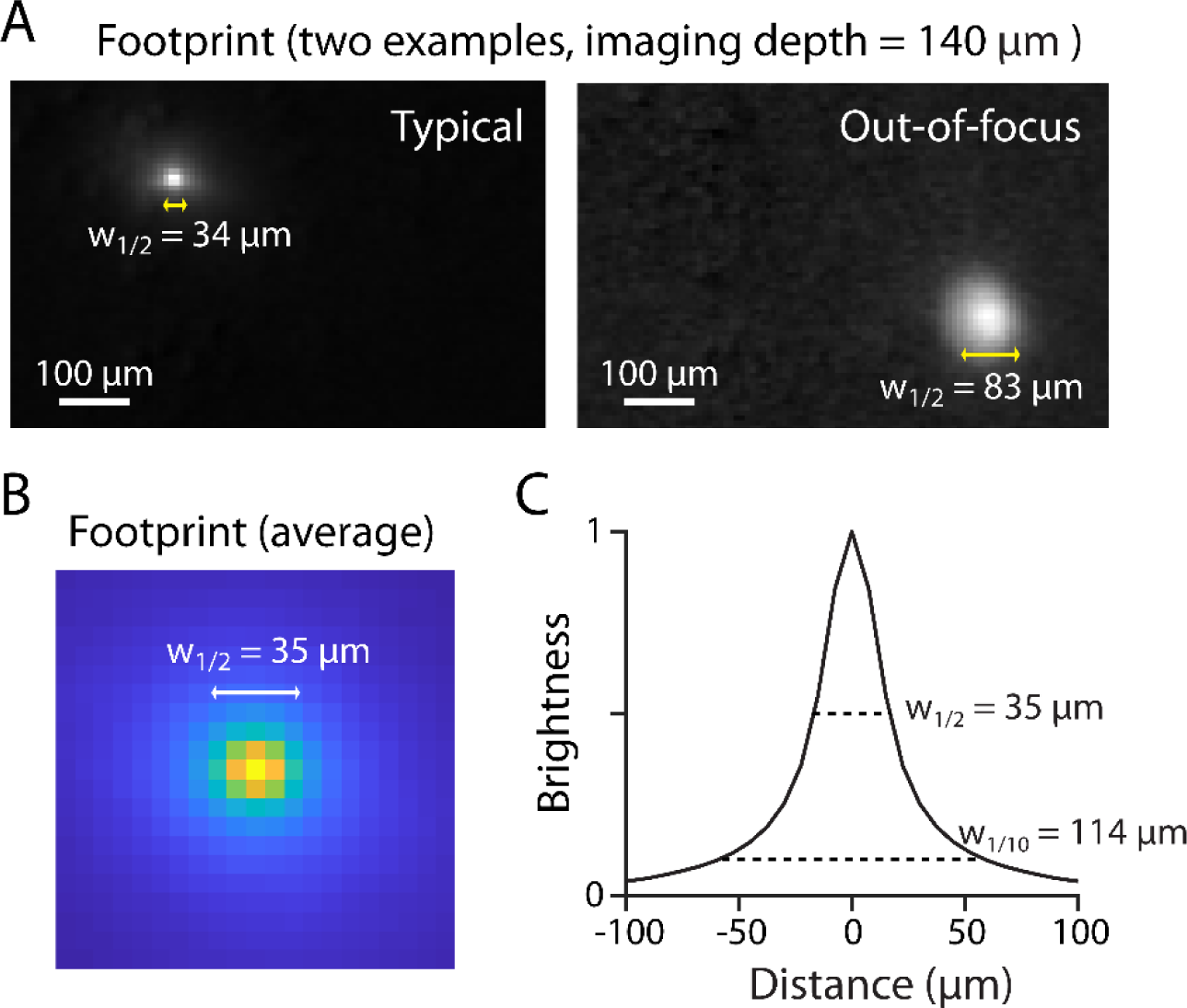
Spatial extent of neuronal footprints in vivo (A) Two example footprints (imaging depth: 140 μm from the surface of CA1), measured by averaging ΔF-AP images of the same cluster. The left image shows the footprint of a typical in-focus neuron, with a full width (w_1/2_) of ∼34 μm at 50% the peak intensity. The right image shows an out-of-focus footprint, with a much wider width (w_1/2_ = 83 μm) **(B)** Footprints aligned to their peaks, and averaged for all cells. **(C)** Normalized brightness of the averaged footprint vs. distance from center. Numbers on the right show the width at 50% (w_1/2_) and 10% (w_1/10_) peak intensity.

**Fig. S5.**
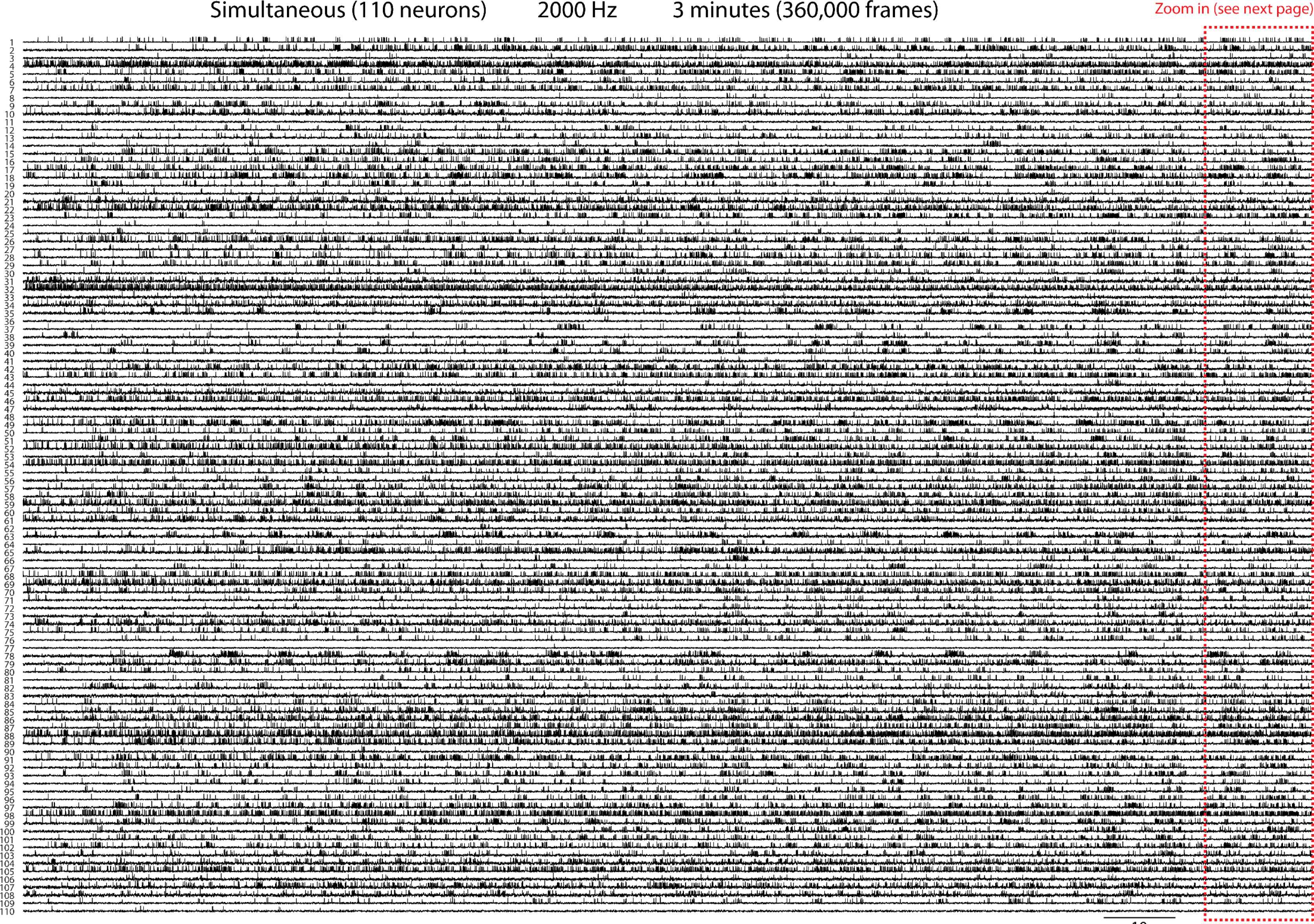
Population spiking dynamics over 3-minute period. Concurrent spike trains of 110 neurons over a 3-minute period, imaged at 2000 frames per second (360,000 frames). Each row shows the extracted activity trace of one neuron. The region encircled by the red box is shown at an expanded scale in Fig. S5.

**Fig. S6.**
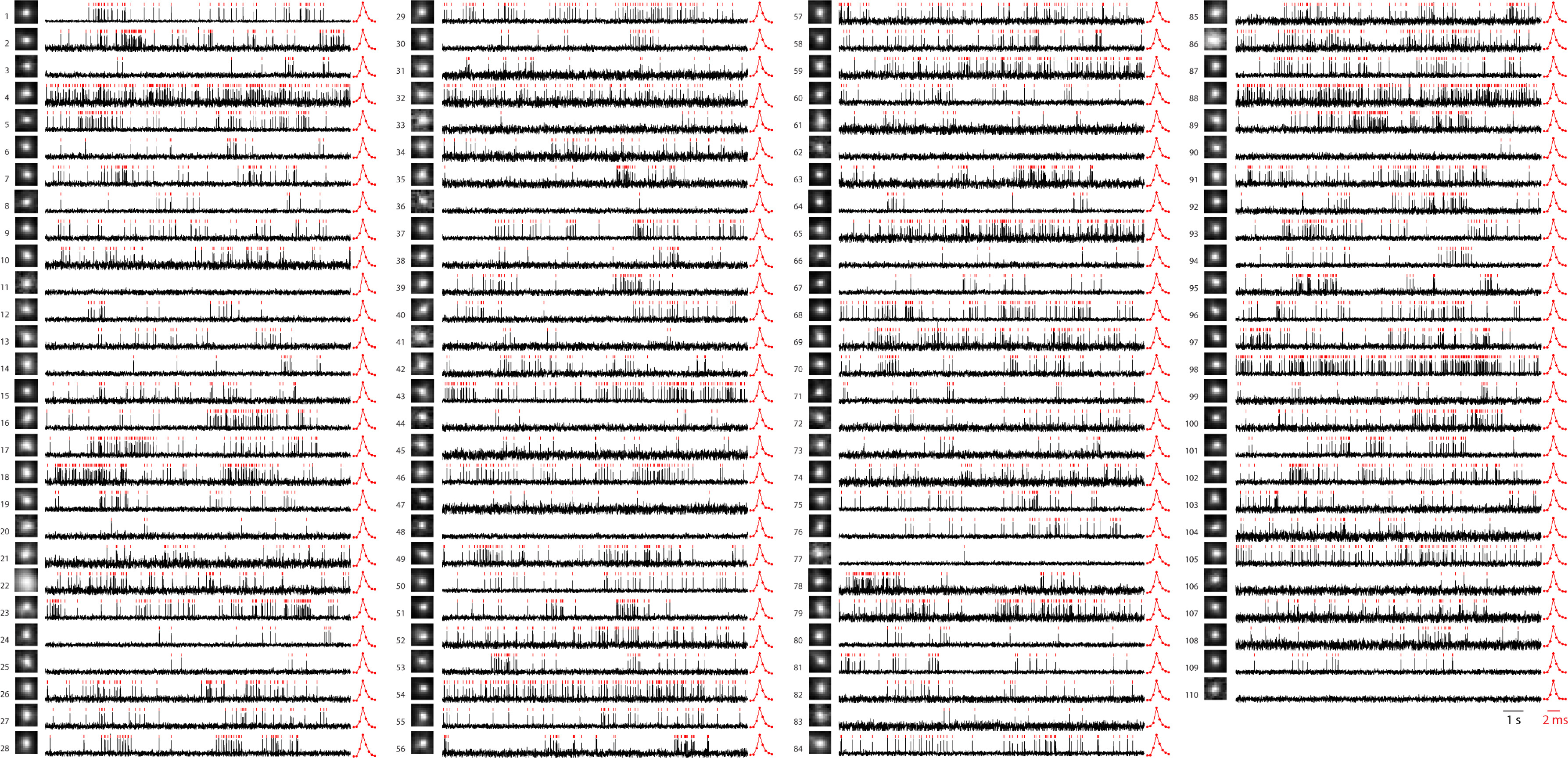
Spike patterns, footprints, and waveforms at higher resolution. Spike patterns of the same 110 neurons in Fig. S4, during the last 15 s of the recording, shown at an expanded scale. Each row shows the activity trace of one neuron. The red ticks above each trace show the detected APs. The image to the left of each trace shows the corresponding footprint (image dimension: 82.5 μm × 82.5 μm). The red trace on the right shows the averaged single-AP waveform of each cell.

**Fig. S7.**
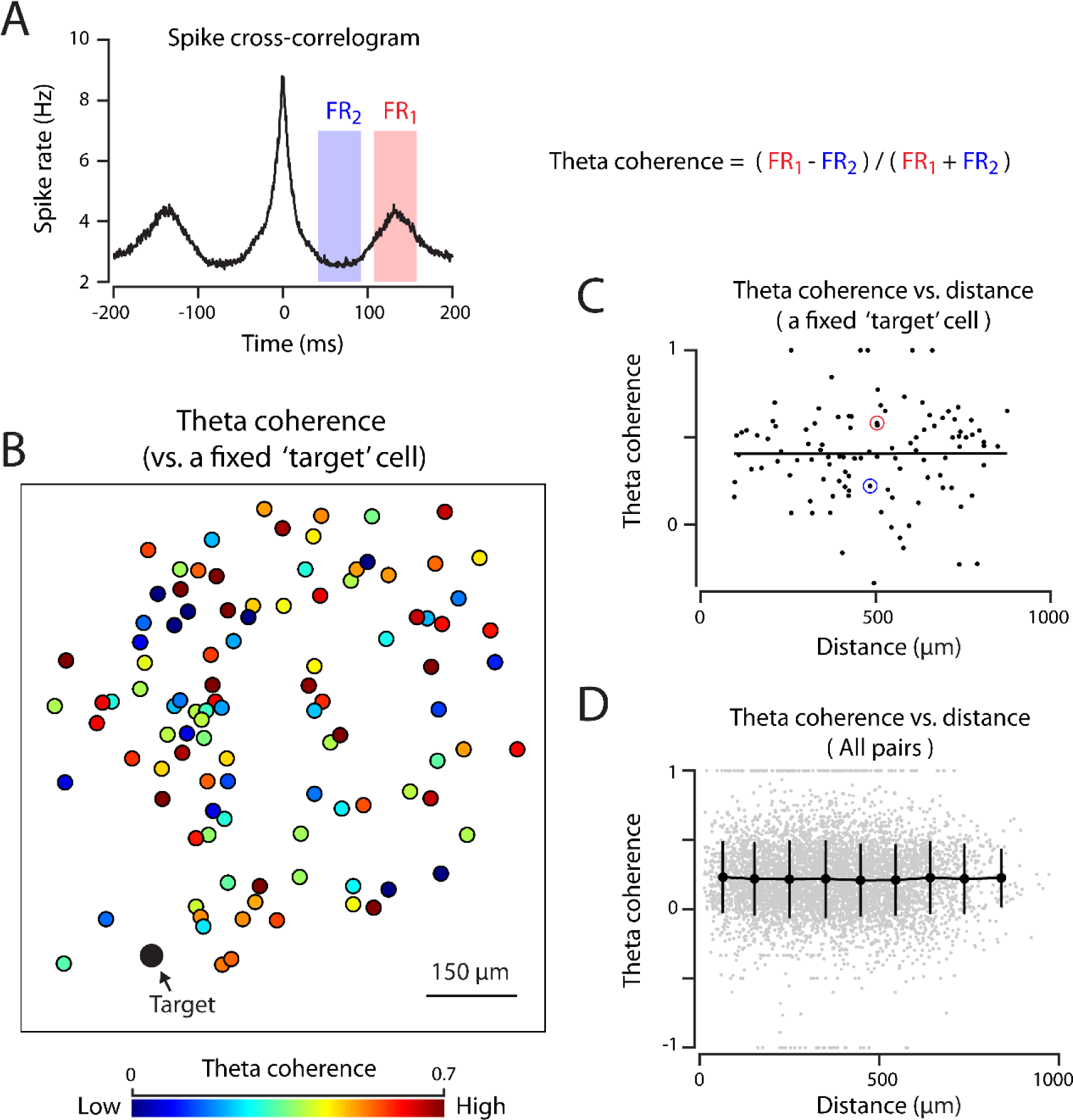
Spatial organization of theta coherence. (A) Definition of theta coherence. For each pair, we measured the mean spike rate of the target cell in two specific time windows (marked by red and blue colors, respectively). Here, FR1 and FR2 represent the target spike rate averaged over the window marked by red and blue colors, respectively. The location of these windows was chosen to match the peak and trough of the population-averaged cross-correlogram (the black trace) at theta frequency. The theta coherence of each cell pair was defined as (FR1-FR2)/(FR1+FR2). (B) A map of theta coherence to a specific target cell (indicated by the black arrow). Each circle shows a neuron, with colors representing the theta coherence relative to the target. (C) Theta coherence (vs. the target cell in B) plotted against the distance from target. The red and blue circles mark the two reference neurons shown in the main figure (red: ref 1, blue: ref 2). The black line shows the least square fit. (D) The coherence between all pairs of neurons plotted against distance. The black curve shows mean ± SD within each distance bin.

**Movie S1.**
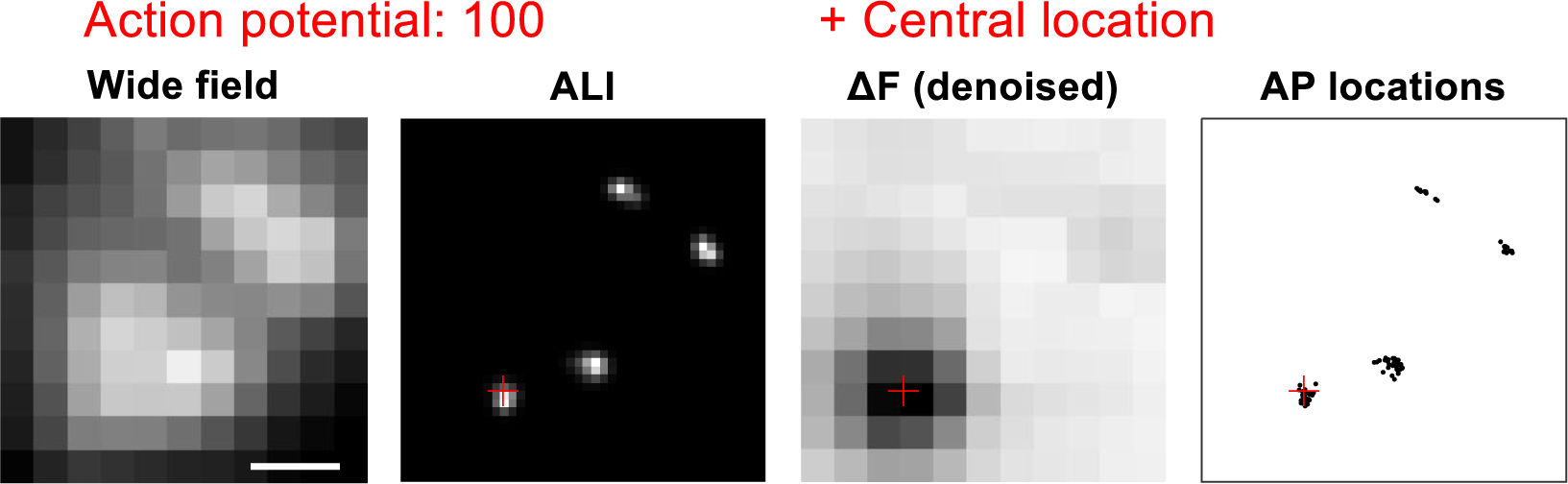
Construction of the ALI map. This movie shows the process of constructing an ALI image from the locations of 100 action potential events. From left to right, wide field image, ALI image, ΔF images of individual AP events (denoised), and the central locations of all APs. The red crosses indicate the central location of the current AP event.

**Movie S2:**
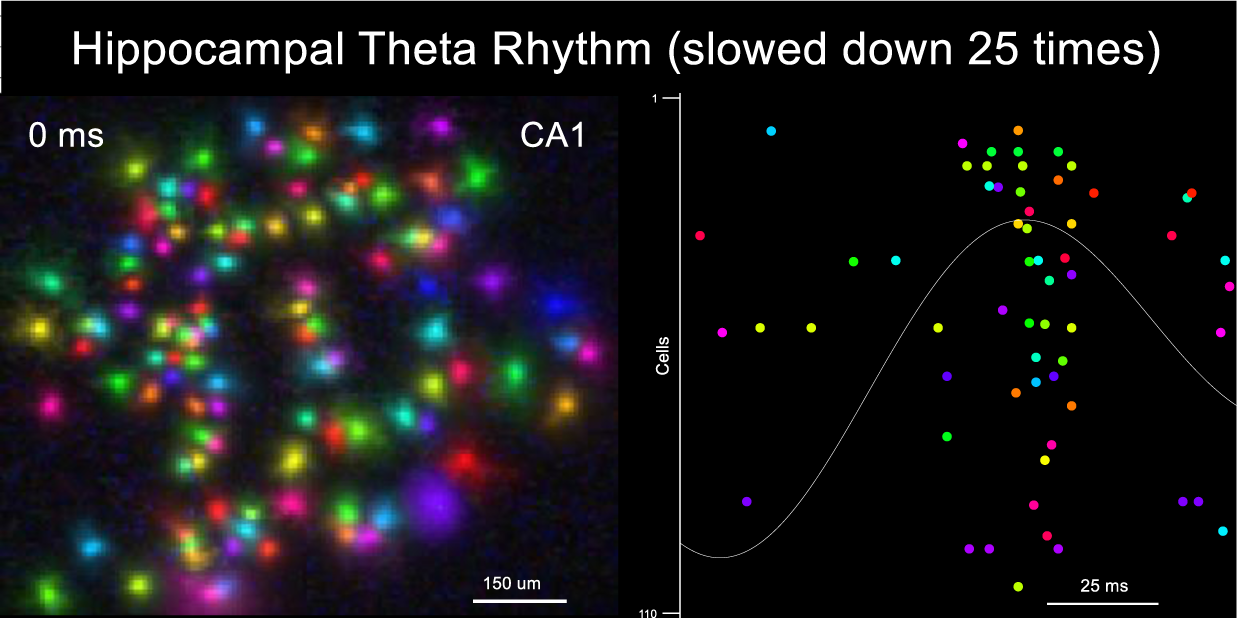
Kilohertz maps of population activity in CA1. This movie shows one second of CA1 ensemble spiking dynamics, slowed down 25 times. Each neuron is shown by a different color. The spatial location of active neurons is mapped at 0.5 ms temporal resolution and shown as a spatial map in the left. The ensemble spiking patterns vs. time are shown in the right. The white curve shows the population theta rhythm, representing the summed activity across neurons and then filtered at the theta-frequency band.

